# Endocytic Patch Dynamics are Differentially Regulated at Distinct Cell Sites in Fission Yeast

**DOI:** 10.1101/2024.12.22.630005

**Authors:** Bethany F. Campbell, Uma J. Patel, Ashlei R. Williams, Maitreyi E. Das

## Abstract

Endocytosis promotes polarity and growth in eukaryotes. In *Schizosaccharomyces pombe* fission yeast, endocytosis occurs at the polarized cell ends and division site and at the non-polarized cell sides. Our characterization of endocytic actin patches show that they are differentially regulated. The patches at the cell ends and division site internalize successfully while those at the sides are weak and erratic. The major regulator of cell polarity, Cdc42, and its target Pak1 kinase only localize to the cell ends and division site. We find that these proteins regulate assembly and internalization of patches at these sites but not at the cell sides. Moreover, Cdc42 specifically activated by the GEF Gef1 promotes proper patch dynamics. Endocytosis requires phosphorylation of the Type I Myosin Myo1 by the Pak1 kinase. Myo1 localizes to the cell ends, division site, and the cell sides. We find that unlike Cdc42 and Pak1, Myo1 also promotes patch assembly at the cell sides. Our data indicate that while Myo1 can globally promote branched actin assembly, successful endocytic patch dynamics and internalization at polarized sites require Cdc42 and Pak1 kinase.

**SUMMARY STATEMENT:** Endocytic patch dynamics are differentially regulated at distinct sites such as the cell ends, division site and the cell sides by Cdc42 and its downstream targets Pak1 kinase and the Type 1 myosin.

## INTRODUCTION

Endocytosis is an essential process by which cells uptake nutrients and recycle proteins from the plasma membrane. In cell-walled organisms such as the fission yeast *Schizosaccharomyces pombe*, internal turgor pressure is high at ~10 atm (Basu et al., 2014; Lacy et al., 2018), roughly equivalent to a racing bike tire. To overcome such high turgor pressure, successful clathrin-mediated endocytosis in yeasts requires synthesis of branched actin networks to produce enough counteractive force to invaginate the plasma membrane and subsequently internalize endocytic vesicles (Aghamohammadzadeh and Ayscough, 2009; Basu et al., 2014; Nickaeen et al., 2022). In *S. pombe*, branched actin is formed by the Arp2/3 complex with the help of nucleation promoting factors such as the Type I myosin Myo1 and the WASP Wsp1 (Sirotkin et al., 2005). Actin binding proteins such as fimbrin Fim1 also localize to endocytic patches to crosslink the branched actin and stimulate force production within the patch (Skau et al., 2011). Furthermore, if endocytosis is impaired, cells cannot grow or divide. Once treated with the Arp2/3 complex inhibitor, CK-666, *S. pombe* cells halt their growth during interphase and fail to form a septum during cytokinesis (Onwubiko et al., 2019). Thus, regulation of branched actin formation is an essential cellular process for *S. pombe* cell viability.

In rod-shaped *S. pombe* cells, clathrin-mediated endocytic events primarily occur at sites of polarization, namely the cell ends during interphase and the division site during cytokinesis (Gachet and Hyams, 2005; Sirotkin et al., 2010). However, some endocytic actin patches also form at the non-growing cell sides. Previous work has shown that sites of endocytosis coincide with regions of cell growth and cytokinesis (Gachet and Hyams, 2005). The GTPase Cdc42 is the major regulator of polarity in *S. pombe* and is active only at polarized cell ends and the division site and not at the cell sides (Das et al., 2012). Additionally, Cdc42 is activated at the division site during actomyosin ring formation and persists throughout cytokinesis (Hercyk and Das, 2019a; Wei et al., 2016). Further work has connected Cdc42 activation to Arp2/3-mediated endocytosis since both are required for septum formation and onset of actomyosin ring constriction in fission yeast (Campbell et al., 2022; Onwubiko et al., 2019). Studies in mammalian MDCK cells also indicate that Cdc42 regulates both endocytic trafficking and exocytic secretion (Kroschewski et al., 1999). Specifically, expression of dominant negative Cdc42^T17N^ mislocalizes basolateral membrane proteins to the apical membrane, while expression of constitutively active Cdc42Q61L depolarizes actin from the apical membrane, causing actin to instead appear throughout the cell periphery (Kroschewski et al., 1999). In *S. pombe*, Cdc42 is independently activated by two GEFs, a transient and mostly cytoplasmic GEF, Gef1, which localizes to cortical puncta at polarized regions (Coll et al., 2003; Das et al., 2015; Hercyk and Das, 2019b), as well as a strong GEF, Scd1, which localizes throughout polarized regions and promotes sustained Cdc42 activation via a positive feedback loop (Das et al., 2012; Das and Verde, 2013; Lamas et al., 2020). Like Cdc42, both of these GEFs are present at polarized cell ends, but it remains unclear which one arrives first at this site to initiate Cdc42 activation, although Scd1 has been shown to prevent ectopic localization of Gef1 (Hercyk et al., 2019). Previous research has shown that Gef1 is recruited to cortical patches via the F-BAR protein Cdc15 (Hercyk and Das, 2019a; Hercyk and Das, 2019b), which promotes endocytosis and is itself recruited to endocytic actin patches via the endocytic Type I myosin, Myo1 (Arasada and Pollard, 2011; Carnahan and Gould, 2003). Conversely, Cdc15 does not recruit Scd1 to cortical patches (Hercyk and Das, 2019b). Additionally, loss of *gef1* leads to excessive accumulation and prolonged lifetimes of Cdc15 within endocytic patches, which can be rescued by expression of a constitutively active *cdc42G12V* allele (Adams et al., 1990). Together, these reports suggest that Cdc15 recruits Gef1 to endocytic patches to promote Cdc42 activation and that this recruitment is important to regulate endocytic patch dynamics.

Prior research has additionally linked Cdc42 activity to the regulation of endocytosis via its direct downstream effector, the p21-activated kinase (PAK) (Murray and Johnson, 2001; Ottilie et al., 1995; Wu et al., 1997), which is activated by binding GTP-bound Cdc42 to relieve its autoinhibition (Bokoch, 2003). Like Cdc42, PAK is conserved across eukaryotic life from yeast to humans, functioning as a serine/threonine kinase (Manser et al., 1994; Marcus et al., 1995). In unicellular eukaryotes, PAK is known to phosphorylate the endocytic Type I myosin in organisms such as slime molds, amoebas, and yeasts (Brzeska et al., 1997; Lechler et al., 2001; Lechler et al., 2000; Lee et al., 2000; Murray and Johnson, 2001; Wu et al., 1997), including *S. pombe* (Attanapola et al., 2009). In these organisms, PAK promotes the ATPase activity and motor function of the Type I myosin by phosphorylating a conserved serine or threonine residue located in the motor domain known as the TEDS-site (Bement and Mooseker, 1995; Fujita-Becker et al., 2005; Pedersen et al., 2023). Specifically, TEDS-site phosphorylation facilitates association of the myosin with actin, stabilizing the complex (Fujita-Becker et al., 2005). In *S. pombe*, phospho-null mutation of the TEDS-site abrogates proper localization of Myo1 to endocytic actin patches, instead causing the myosin to localize uniformly along the plasma membrane (Attanapola et al., 2009). Additionally, we have recently shown that Pak1 promotes successful internalization of endocytic patches and even its own removal via endocytosis at growing cell ends of *S. pombe* (Harrell et al., 2024). These findings suggest that regulation of Cdc42 activation may be necessary to facilitate proper endocytic patch dynamics.

While we see distinct sites of endocytosis at the cells ends, division site, and the cell sides, it is not known whether these sites are functionally similar and if they show different regulatory patterns. Here, we use genetic approaches and quantitative live cell imaging to better define the intricate connections between the different sites of polarization and endocytosis. We show that endocytic actin patches display disparate dynamics at distinct cell sites. Endocytosis at the polarized sites requires Cdc42, its target Pak1, and the downstream substrate Myo1. At the cell sides, Myo1 but not Cdc42 and Pak1 regulates patch assembly and dynamics. While our recent work shows that endocytosis ensures proper spatiotemporal regulation of Cdc42 activation (Harrell et al., 2024), here we demonstrate that Cdc42 activation likewise spatiotemporally directs endocytosis.

## RESULTS

### Endocytic events at distinct regions in the cell show differential dynamics

To our knowledge, the dynamic behaviors of individual endocytic patches within each distinct region have not been characterized and reported. We were thus curious to determine how endocytic patches at polarized regions behave compared to those at non-growing regions. To do this, we performed 1-sec interval timelapse imaging of asynchronous cultures of wild-type cells expressing mEGFP-tagged fimbrin Fim1, a branched actin crosslinker (Nakano et al., 2001), which promotes actin bundling necessary for endocytic patch internalization (Skau et al., 2011). This approach allowed us to simultaneously capture the dynamics of individual endocytic patches at the cell ends, the cell sides, and the division site (Fig. 1 A). We then used the FIJI plugin TrackMate (Tinevez et al., 2017) to track the trajectories of individual Fim1-mEGFP patches. From this data, we found that Fim1-mEGFP-labeled endocytic patches have disparate lifetimes at each region, with patches at the cell sides exhibiting the shortest lifetimes, while patches at the division site show the longest lifetimes (Fig. 1 B). While the lifetimes at polarized cell ends and the division site are statistically different in duration, the mean lifetimes for both regions (~16 sec at the cell ends and ~18 sec at the division site) fall within the reported average range for Fim1-mEGFP lifetimes (Berro and Pollard, 2014; Sirotkin et al., 2010). In contrast, the patch lifetime at the cell sides is ~12 sec (Fig. 1 B), well below than that at the polarized sites.

**Figure. 1.**
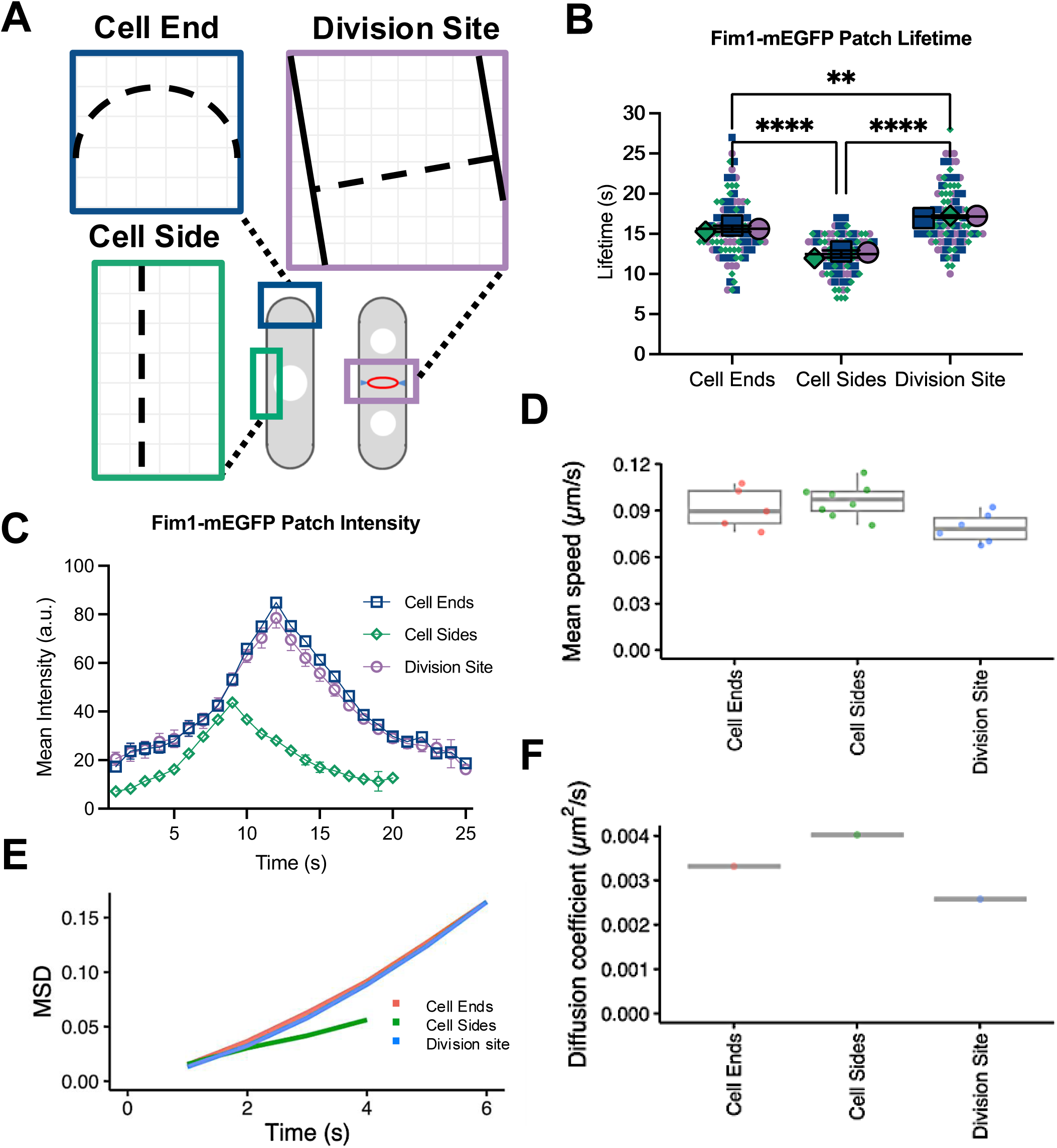
Endocytic events at distinct sites in the cell show differential dynamics. **A.** Endocytic events occur at different regions of polarized *S. pombe* cells. Colored tracks indicate trajectories of individual Fim1-mEGFP patches. **B.** Lifetimes of Fim1-mEGFP labeled endocytic patches at each region. **C.** Mean Fim1-mEGFP peak intensity in endocytic patches at each region. **D.** Mean speed of endocytic patches at each region averaged from individual cells. **E.** MSD (mean square displacement) of Fim1-mEGFP patches calculated for each region. **F.** Diffusion coefficient of Fim1-mEGFP patches at each region. (n ≥ 28 patches per region, N=3; n ≥ 6 cells analyzed per region). Ordinary one-way ANOVA with Tukey’s multiple comparisons. ****, *p*< 0.0001; **, *p*< 0.01

Mean patch intensity was also measured frame-by-frame throughout each patch’s lifetime. This analysis revealed that endocytic patches at the cell ends and the division site recruit Fim1-mEGFP at the same rate and to the same peak intensity (Fig. 1 C). Conversely, endocytic patches at the cell sides recruited ~50% less Fim1-mEGFP at peak intensity compared to the polarized sites (Fig. 1 C). Using the R package TrackMateR (Sittewelle and Royle, 2024), we calculated the diffusive properties of tracked Fim1-mEGFP patches to characterize their respective dynamics. Calculations show that endocytic patches at the division site are the slowest moving (Fig. 1 D) and most stable with the least diffusion of all sites (Fig. 1 F). In contrast, endocytic patches at the cell sides move the fastest (Fig. 1 D) and exhibit the highest diffusion rate (Fig. 1 F). Patches at the cell ends display a comparatively intermediate speed and diffusion rate (Fig. 1 D and F). However, mean squared displacement (MSD) analysis reveals that endocytic patches at the cell ends and the division site exhibit similar degrees of directed motion, while patches at the cell sides demonstrate more diffusive behavior overall (Fig. 1 E). Together, these data indicate that endocytic patches at polarized regions recruit twice as much Fim1-mEGFP, are longer-lived, and exhibit more stable and directed dynamics compared to patches at the non-growing cell sides.

### Cdc42 activity promotes timely formation of endocytic branched actin networks at the division site

Our findings that endocytic events behave differently at polarized regions compared to non-growing cell sides suggest that polarity regulators may influence endocytic dynamics. Indeed, Cdc42 has been shown to promote uptake of Lucifer Yellow via endocytosis in *S. pombe* (Murray and Johnson, 2001). Connecting previous observations that sites of endocytosis mirror sites of Cdc42 activation, we asked if Cdc42 activation is necessary for endocytosis. To test this idea, we asked if a delay in Cdc42 activation at the division site would consequently delay onset of endocytic activity within the division plane. We selected the division site since the temporal pattern of Cdc42 activation is best understood at this site. The GEF Gef1 is the first GEF to appear and activate Cdc42 at the division site, and loss of Gef1 consequently leads to a delay in Cdc42 activity at this site (Wei et al., 2016) since the other Cdc42 GEF, Scd1, depends on Gef1 for timely localization to the division site (Hercyk et al., 2019). While delayed Cdc42 activation in *gef1Δ* mutant delays actomyosin ring constriction due to delayed recruitment of Bgs1, the timing of mitotic events proceeds normally (Wei et al., 2016). Thus, we measured the timing of Fim1-mEGFP appearance at the division site relative to onset of mitosis marked by spindle pole body (SPB) separation in *gef1Δ* cells compared to *gef1^+^* controls. These experiments show that Fim1-mEGFP first appears at the division site ~12 mins after SPB separation in *gef1^+^* cells, while its appearance in *gef1Δ* cells is delayed to ~20 mins post-SPB separation (green dotted box, Fig. 2A and Supplementary Fig.S1A). Intriguingly, Cdc42 is also first activated at the division site ~10-12 min post-SPB separation in wild-type cells (Hercyk and Das, 2019b; Wei et al., 2016). Fim1-mEGFP patches appear uniformly throughout the division site ~20 mins post-SPB separation in *gef1^+^* controls compared to ~28 mins post-SPB separation in *gef1Δ* mutants (solid green box, Fig. 2 A and B). These findings suggest that Cdc42 activation is indeed required for endocytosis.

**Figure 2.**
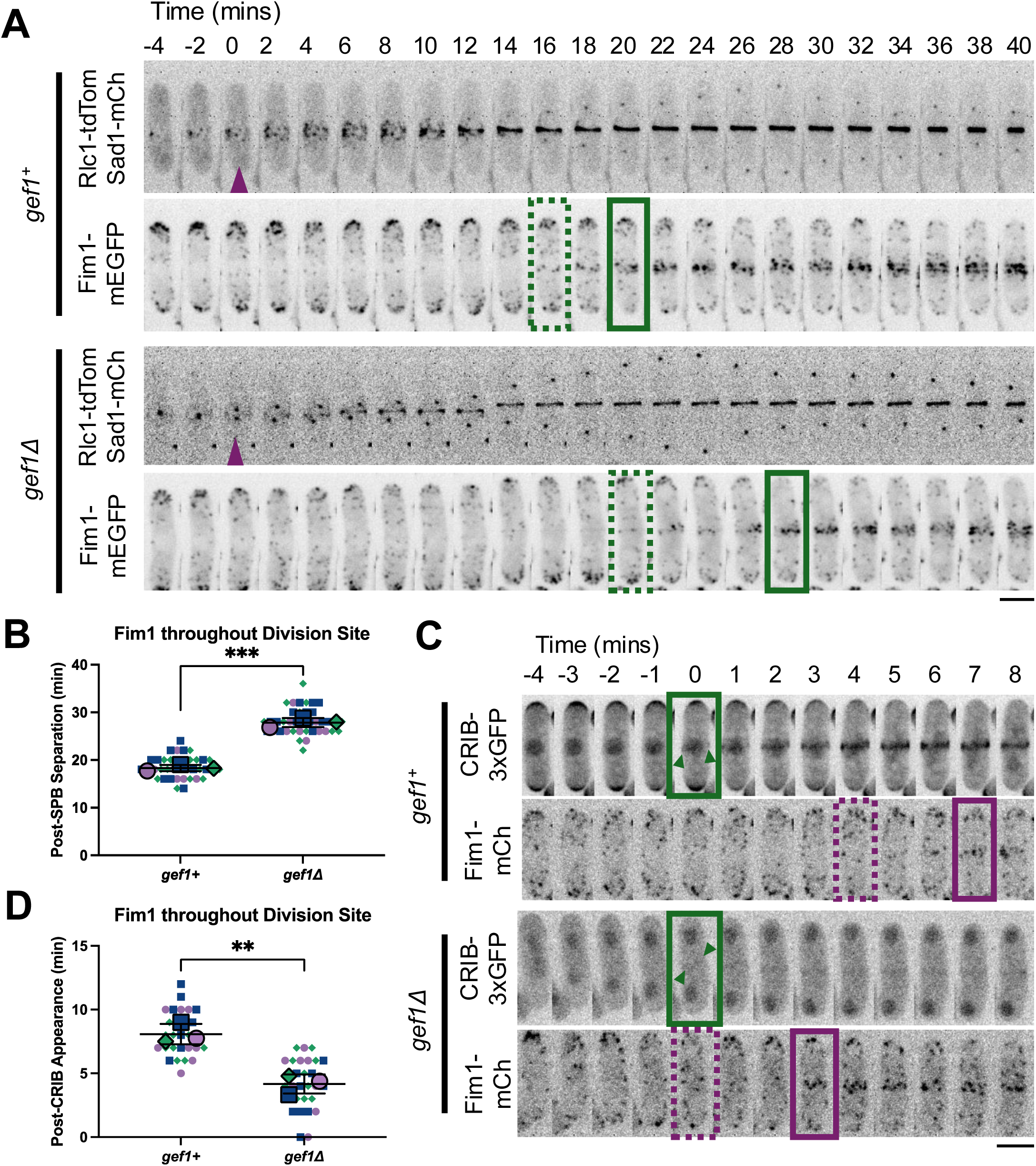
Endocytosis follows Cdc42 activation at the division site. **A.** Representative cells showing initial (green dotted box) and uniform (solid green box) Fim1-mEGFP appearance at the division site relative to Sad1-mCherry labeled spindle pole body (SPB) separation (purple arrowhead). **B.** Quantification of A, where time 0 = SPB separation. **C.** Representative cells showing initial (purple dotted box) and uniform (solid purple box) Fim1-mCherry appearance relative to CRIB-3xGFP appearance (green box) at the division site. **D.** Quantification of C, where time 0 marks CRIB-3xGFP appearance (green arrowheads) at the division site. (n ≥ 8 cells per genotype per experiment). Unpaired Student’s *t*-test. ****, *p*< 0.0001; ***, *p*<0.001; **, *p*<0.01, scale bar=5μm

We were then curious to determine how long after Cdc42 activation does Fim1 appear at the division site in *gef1^+^* as well as *gef1Δ* cells. In *gef1^+^* cells, Fim1-mCherry first appears at the division site (purple dotted box) ~5-6 mins after Cdc42 is activated (green box and arrowheads, Fig. 2 D and Supplementary Fig.S1B), as visualized by the CRIB-3xGFP bio-probe which binds active GTP-bound Cdc42 (Tatebe et al., 2008). Comparatively, in *gef1Δ* cells, Fim1-mEGFP first appears at the division site (purple dotted box) about the same time as Cdc42 activation (green box and arrowheads, Fig. 2 D and Supplementary Fig.S1B). Similarly, Fim1-mCherry appears throughout the division site after Cdc42 activation at ~7-8 mins in *gef1^+^* and at ~4 mins in *gef1τι* cells (Fig. 2 C and D). This suggests that endocytosis starts soon after Cdc42 activation in *gef1Δ* mutants since these cells have already progressed through mitosis and thus many cellular events are already complete. Our findings suggest that *gef1Δ* cells simply await Cdc42 activation to initiate endocytosis. Together, these data suggest that Cdc42 activation is required for the onset of endocytosis at the division site.

### The GEFs Gef1 and Scd1 differentially regulate Cdc42 to direct branched actin synthesis and dynamics at sites of polarization

While the Cdc42 GEF Scd1 is required for polarized growth at the cell ends, Gef1 is required for bipolar growth (Hercyk et al., 2019). Findings that Gef1 and Scd1 differentially activate Cdc42 at polarized sites prompted us to define the manner of Cdc42 activation required to direct endocytosis. We tracked the dynamics of Fim1-mEGFP at the ends in *gef1Δ* and *scd1Δ* cells compared to *gef1^+^scd1^+^* controls, as previously described (Fig. 1). In these experiments, cells also expressed Cdc15-tdTomato to corroborate and expand upon our previous findings. As shown before (Onwubiko et al., 2019), *gef1Δ* mutants exhibit excessive accumulation and prolonged lifetimes of Cdc15-tdTomato at the cell ends compared to *gef1^+^* control cells (Fig. 3 A; Supplementary Fig. S2 A and B). We additionally find that Fim1-mEGFP accumulation within endocytic patches is likewise increased in *gef1Δ* mutants at both the cell ends and the division site compared to controls. (Fig. 3 A, B, and D). In in *gef1Δ* mutants, Fim1-mEGFP lifetimes are also prolonged at the cell ends; however, we saw no change in patch lifetime at the division site in *gef1Δ* cells (Fig. 3 H). Notably, the recruitment patterns and lifetimes of Fim1-mEGFP patches are the same at the cell sides of *gef1^+^* and *gef1Δ* cells (Fig. 3 F and H), which suggests that the dynamics of these patches are not influenced by Cdc42 or its GEFs.

**Figure 3.**
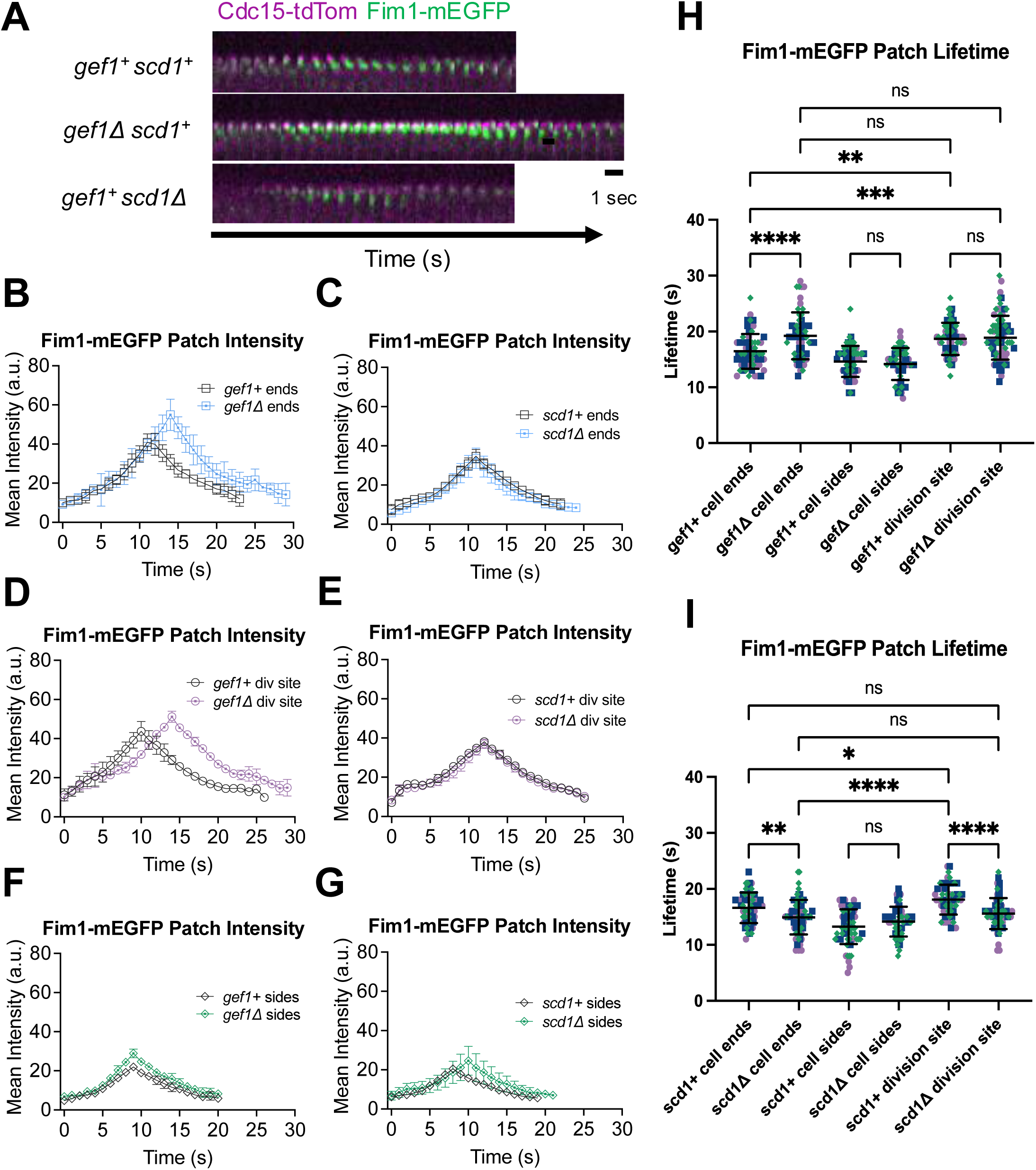
The Cdc42 GEFs Gef1 and Scd1 differentially regulate endocytosis. **A.** Representative montages of Cdc15-tdTom and Fim1-mEGFP at growing cell ends in the indicated genotypes. **B-G.** Mean patch intensities of Fim1-mEGFP at growing cell ends, the division site, and cell sides. (n ≥ 20 endocytic patches per triplicate experiment). **H.** and **I.** Patch lifetimes of Fim1-mEGFP at each region in the indicated genotypes. (n ≥ 20 endocytic patches, N=3). Ordinary one-way ANOVA with Tukey’s multiple comparisons. ****, *p*<0.0001; ***, *p*<0.001; **, *p*<0.01; *, *p*< 0.05; ns, not significant

In contrast, in *scd1Δ* mutants, we did not observe any change in the recruitment dynamics of either Cdc15-tdTomato or Fim1-mEGFP compared to *scd1^+^* cells (Supplementary Fig. S2 C and D; Fig. 3 A, C, E, and G). We did, however, find that Fim1-mEGFP patches are depolarized rather than concentrated at the cell ends in *scd1Δ* mutants (Supplementary Fig. S2 E). Fim1-mEGFP patches in *scd1Δ* mutants also display similar lifetimes at all regions of the cell (Fig. 3 H), which is consistent with the random, depolarized Cdc42 activation patterns that occur in these cells due to loss of Scd1-mediated restriction of ectopic Gef1 localization (Hercyk et al., 2019). Together, these results suggest that Gef1 is the primary GEF that regulates Cdc42 activation to direct proper endocytic patch dynamics at polarized regions.

### Pak1/Orb2 kinase activity promotes timely formation of endocytic branched actin networks at the division site

Next, we asked if Cdc42 regulates endocytosis via its target the Pak1 kinase. In addition to the cell ends, Pak1 is also present at the division site during cytokinesis where it promotes assembly of the contractile actomyosin ring (Magliozzi et al., 2020). Similar to previously described experiments (Fig. 2), we measured the time intervals between SPB separation and appearance of Fim1-mEGFP at the division site in *orb2^+^* cells compared to *orb2-34*, a kinase-dead, temperature-sensitive allele of *pak1* (Das et al., 2012; Verde et al., 1995). Even at permissive temperature, *orb2-34* mutants show minimal kinase activity and display polarity defects compared to wild-type *pak1* (Das et al., 2012; Verde et al., 1995). Thus, experiments were performed under permissive 25°C conditions to minimize pleiotropic effects that may arise under temperature restriction, which causes *orb2-34* cells to become rounded and orb-like (Verde et al., 1995). Under permissive conditions, the *orb2-34* mutants grow as well as *orb2^+^* cells and only display cell morphology defects. Nevertheless, we do see a ~2 min delay in the appearance of Fim1-mEGFP appearance (dotted and solid green boxes) at the division site in *orb2-34* mutants compared to controls (Fig. 4 A-C). While this is not a major delay, it is consistent and significant. It is possible that under complete abrogation of Pak1 kinase activity the difference will be enhanced. However, our experimental conditions do not allow us to expose the cells to such extreme conditions for extended timelapse imaging while avoiding pleiotropic effects. Thus, these data indicate that timely initiation of endocytosis at the division site depends both on Cdc42 and Pak1 activation.

**Figure 4.**
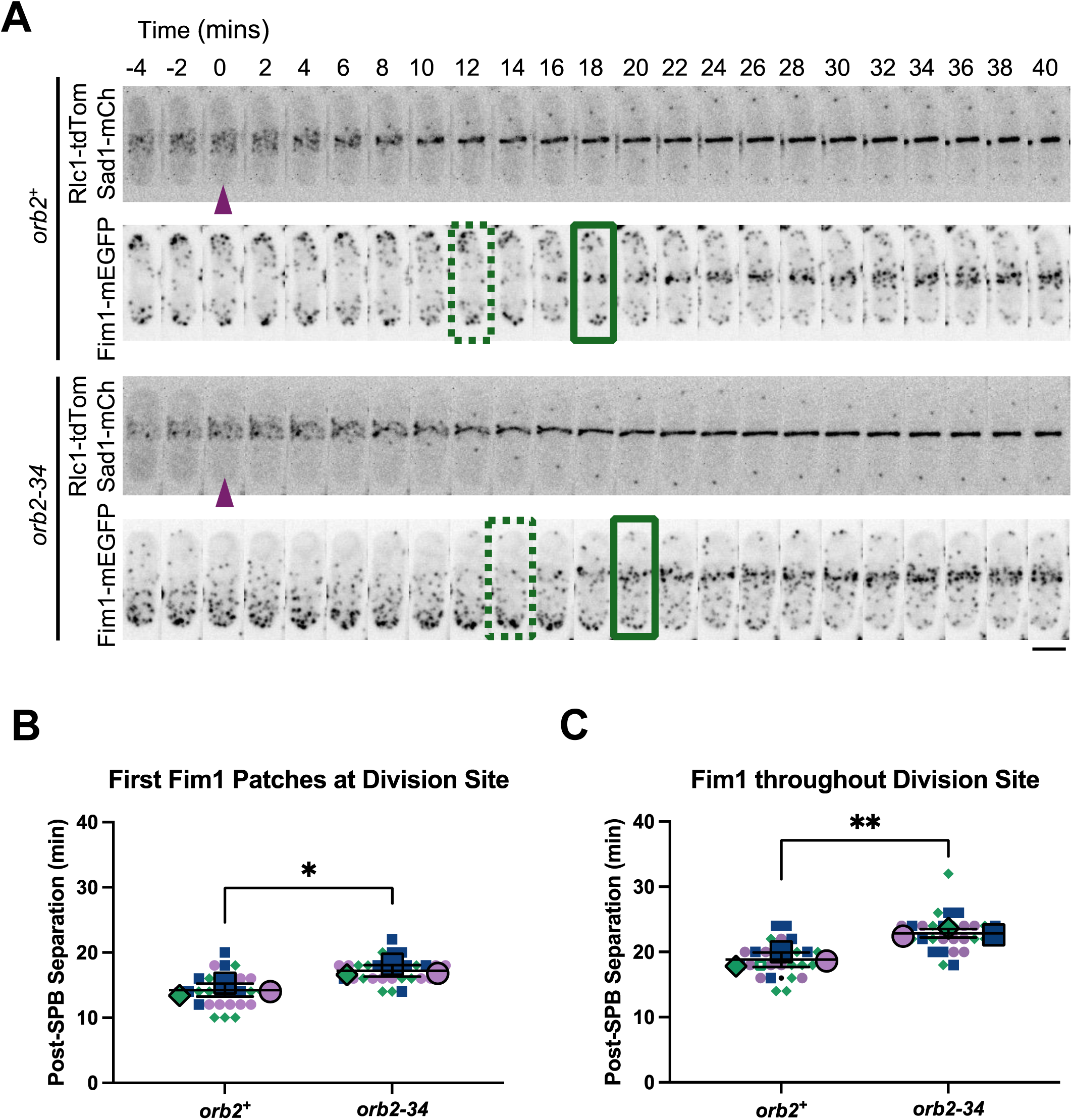
Pak1/Orb2 kinase is required for timely initiation of endocytosis at the division site. **A.** Representative *orb2+* and *orb2-34* cells showing initial (green dotted box) and uniform (solid green box) Fim1-mEGFP appearance at the division site relative to Sad1-mCherry to mark spindle pole body separation (purple arrowhead. **B-C.** Quantification of A, where time 0 = SPB separation. (n ≥ 8 cells per genotype per experiment). Unpaired Student’s *t*-test. **, *p*<0.01; *, *p*< 0.05, scale bar=5μm

### Pak1 facilitates proper formation and dynamics of endocytic actin patches at sites of polarization

Given our recent findings that loss of Pak1 kinase function impairs internalization of endocytic patches at polarized cell ends (Harrell et al., 2024), we asked whether similar endocytic defects also occur at the division site and cell sides. To examine this, we tracked Fim1-mEGFP patches in temperature-sensitive *orb2-34* mutants and compared their dynamics to patches tracked in *orb2^+^* control cells. As at the cell ends (Harrell et al., 2024), Fim1-mEGP patch internalization at the division site is impaired in *orb2-34* mutants compared to controls, especially at the 35°C restrictive temperature (Fig. 5 A). Overall, it appears that Fim1 patches largely move within the plane of the plasma membrane in the absence of Pak1 kinase function.

**Figure 5.**
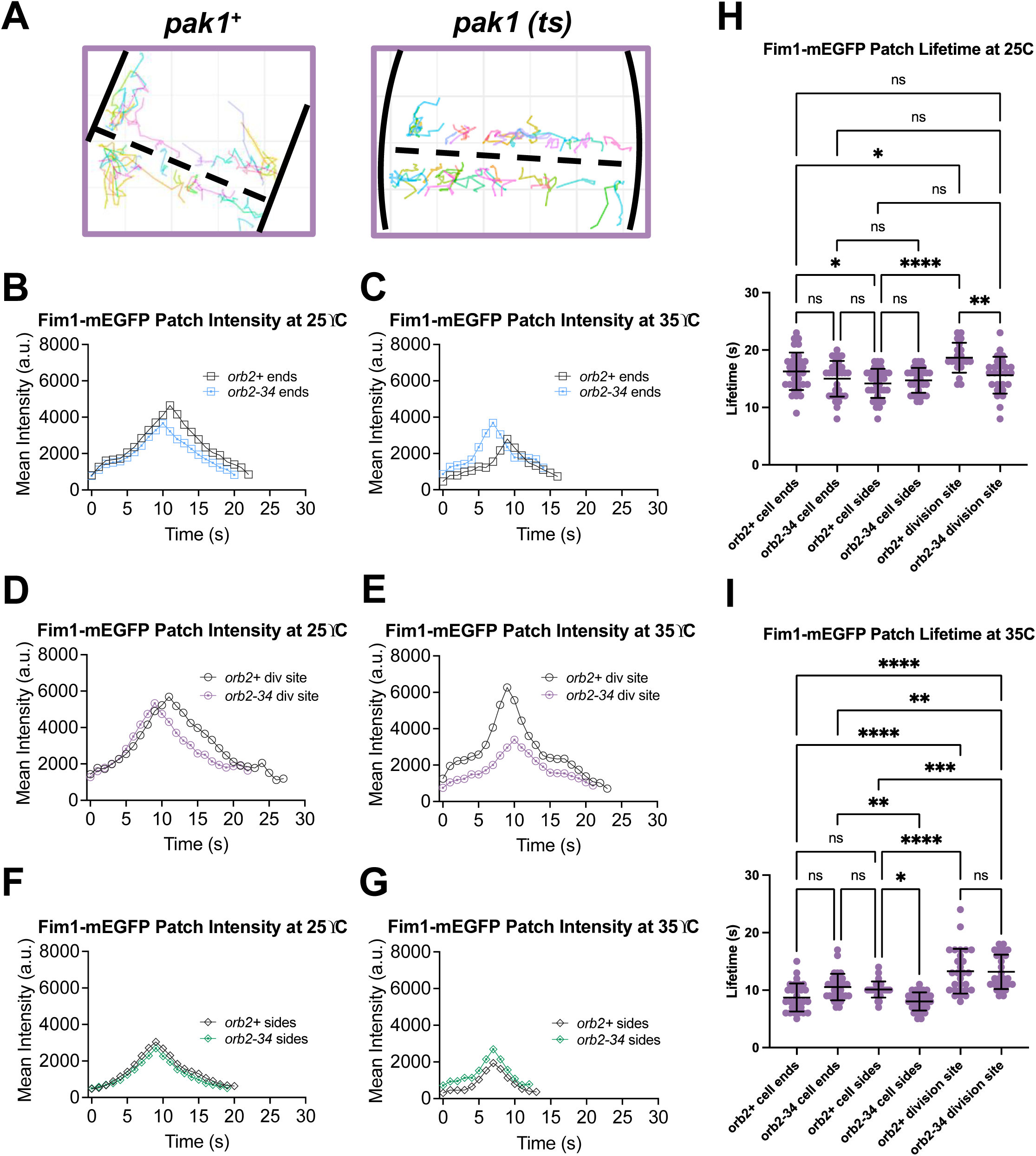
Pak1/Orb2 kinase is required for normal endocytic patch dynamics at the cell ends. **A.** Representative cells showing Fim1-mEGFP endocytic events tracked via TrackMate. **B-G.** Mean patch intensities of Fim1-mEGFP at growing cell ends, the division site, and cell sides in the indicated genotypes and conditions. (n ≥ 20 endocytic patches per experiment). **H-I.** Patch lifetimes of Fim1-mEGFP at each region in the indicated genotypes and conditions. (n ≥ 20 endocytic patches, N=3). Ordinary one-way ANOVA with Tukey’s multiple comparisons. ****, *p*<0.0001; ***, *p*<0.001; **, *p*<0.01; *, *p*< 0.05; ns, not significant

We also observe impaired recruitment of Fim1-mEGFP to endocytic patches in *orb2-34* mutants under permissive conditions. At 25°C, *orb2-34* cells show decreased peak Fim1-mEGFP intensity within endocytic patches at the cell ends and the division site compared to controls (Fig. 5 B and D). In contrast, at the 35°C restrictive temperature, peak Fim1-mEGFP intensity at the cell ends in *orb2-34* mutants remains the same as observed at 25°C, while *orb2^+^* controls show decreased Fim1-mEGFP patch intensity at the cell ends at this temperature (Fig. 5 C). However, we observe the opposite effect at the division site. Here at 35°C, Fim1-mEGFP patch intensity dramatically decreases at the division site in *orb2-34* mutants while patch intensity is unchanged in *orb2^+^* controls (Fig. 5 E). These observations suggest that Fim1 recruitment to the cell ends of *orb2-34* mutants is insensitive to temperature restriction, while Fim1 recruitment to the division site of *orb2-34* mutants is impaired under these conditions. Given that patch internalization is impaired at the cell ends in *orb2-34* mutants (Harrell et al., 2024), our data suggest the level of actin assembly and dynamics alone do not determine the ability of endocytic vesicles to successfully internalize. Consistent with previous experiments showing that endocytic dynamics at the cell sides are not influenced by changes in polarity regulation, we did not observe changes in Fim1-mEGFP recruitment at these sites other than some decrease in peak patch intensity in *orb2^+^* controls at 35°C (Fig. 5 F and G).

Across the board at all cell regions, Fim1-mEGFP patch lifetimes decrease at 35°C compared to 25°C (Fig. 5 H and I), likely due to increased Brownian motion. As previously reported (Harrell et al., 2024), Fim1-mEGFP patch lifetimes do not change at the cell ends of *orb2-34* mutants compared to controls (Fig. 5 H and I). Conversely, at 25°C, Fim1-mEGFP patch lifetimes are ~5 seconds shorter at the division site in *orb2-34* cells compared to *orb2^+^*cells (Fig. 5 H), but this difference is abrogated at 35°C where *orb2^+^* patch lifetimes decrease (Fig. 5 I). While loss of Pak1 kinase function impairs endocytic internalization at both the cell ends and the division site, our data suggests that Pak1 may perform additional functions at the division site since we observe changes in Fim1 accumulation and lifetime at this site, but not at the cell ends. Together, these results indicate that Pak1 kinase plays important and differential roles at polarized regions to regulate endocytic patch behavior.

### The Type I myosin Myo1 promotes timely internalization of endocytic actin patches via Pak1-mediated phosphorylation of its motor domain

Our data show that endocytosis is differentially regulated at distinct sites within polarized cells. While Cdc42 and Pak1 kinase are required for proper actin patch dynamics at the cell ends and the division site, they are not required for the patches at the cell sides. Given that previous reports indicate that the Pak1 kinase regulates endocytosis via Myo1 phosphorylation, we asked if Myo1 itself is differentially required for endocytosis at these distinct sites. We first analyzed Myo1-GFP localization in *S. pombe*. As expected Myo1-GFP localizes to the growing cell ends and the division site (Attanapola et al., 2009; Carnahan and Gould, 2003; Lee et al., 2000). Myo1-GFP puncta also appear along the cell sides, co-localizing with branched actin patches as labeled with Fim1-mCherry (arrowheads, Supplementary Fig. S3A) (Arasada and Pollard, 2011; Sirotkin et al., 2010). Next, we examined Fim1 dynamics in *myo1Δ* cells compared to *myo1^+^*controls at the cell ends, division site, and the cell sides. In the absence of *myo1*, endocytic patches marked with Fim1-mCherry display weak internalization and largely slide within the plane of the plasma membrane at both the cell ends and the division site (Fig. 6 A). Additionally, fewer endocytic patches form in *myo1Δ* mutants, although Fim1-mCherry lifetimes are much longer compared to controls at all regions (Fig. 6 B). Notably, while Fim1-mCherry recruitment to endocytic patches reaches higher peak levels in *myo1Δ* cells compared to *myo1^+^* controls, Fim1 takes longer to accumulate at patches in these mutants (Fig. 6 D). Indeed, peak Fim1-mCherry patch intensity is reached ~8 seconds later at the cell ends (Fig. 6 E) and ~11 seconds later at the division site in *myo1Δ* mutants compared to *myo1^+^* controls due to this slower recruitment of Fim1 (Fig. 6 F). At the cell sides, peak Fim1-mCherry patch intensity is slightly increased in *myo1Δ* cells and patches also take longer to reach peak intensity (Fig. 6 G). Since Myo1 can localize to cell sides in addition to sites of polarization, this may explain the changes in Fim1 behavior at the cell sides in *myo1Δ* mutants.

**Figure 6.**
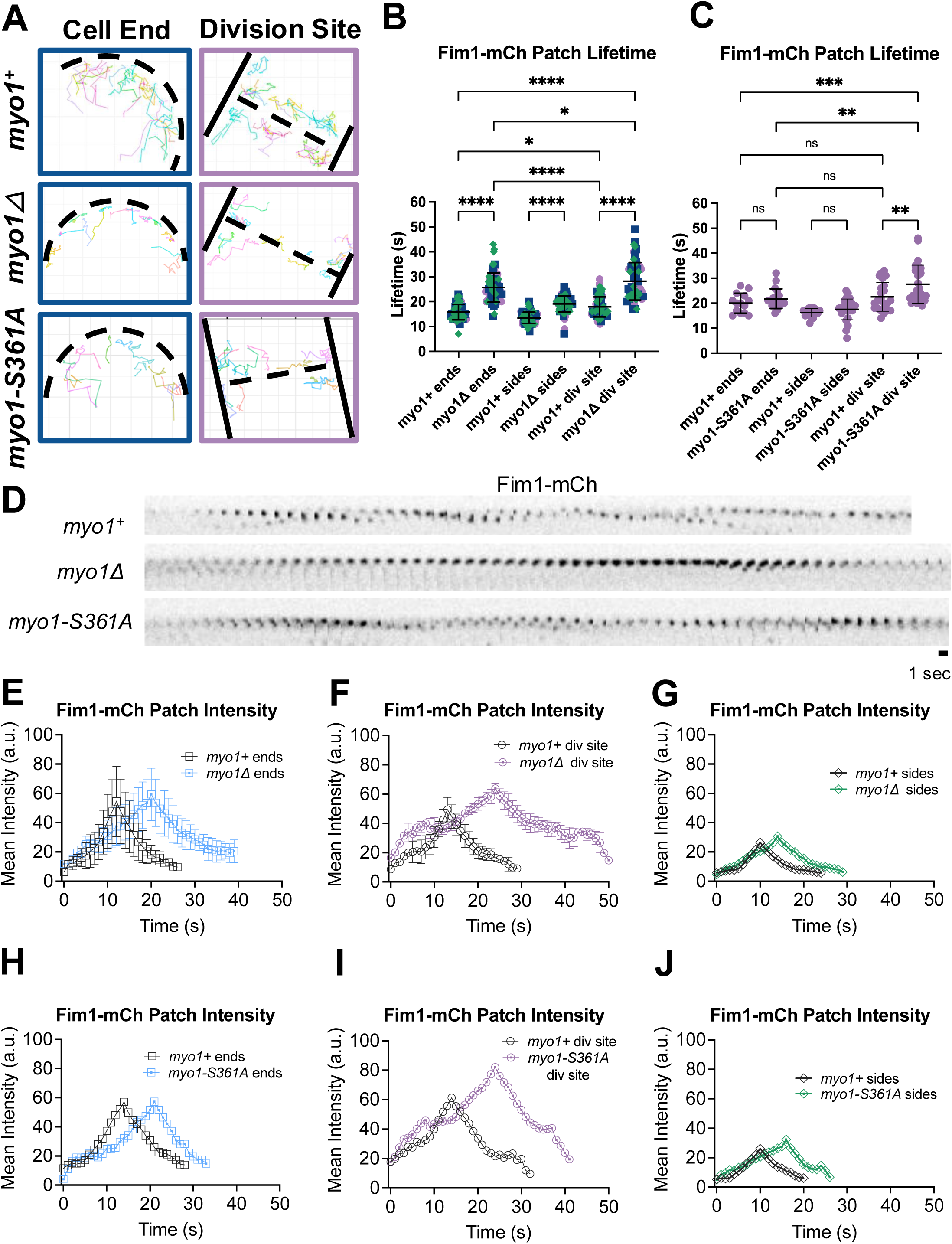
The type 1 Myosin Myo1 regulates endocytic patch dynamics at the cell ends, division site and the cell sides. **A.** Representative cells showing Fim1-mCherry endocytic events tracked via TrackMate in the indicated strains. **B.** Patch lifetimes of Fim1-mCherry at each region in *myo1^+^* and *myo1Δ*. **C**. Patch lifetimes of Fim1-mCherry at each region in *myo1^+^* and *myo1S361A*. **D.** Montages of Fim1-mCherry overtime at the cell ends of the indicated strains. **E-J.** Mean patch intensities of Fim1-mCherry at each region of the indicated strains. (n ≥ 17 endocytic patches, N=3). Ordinary one-way ANOVA with Tukey’s multiple comparisons. ****, *p*<0.0001; ***, *p*<0.001; **, *p*<0.01; *, *p*< 0.05; ns, not significant

While Pak1 phosphorylates Myo1 at the S361 TEDS site located within its N-terminal motor domain (Attanapola et al., 2009), Myo1 also binds and weakly activates the branched actin nucleating Arp2/3 complex via its C-terminal tail (Lee et al., 2000; Sirotkin et al., 2005). Thus, Myo1 may promote endocytic patch internalization via its N-terminal motor function and/or its C-terminal association with Arp2/3. As *myo1Δ* completely abolishes all Myo1 function, experiments in *myo1Δ* mutants alone cannot distinguish between these mechanistic possibilities. To determine if Pak1-mediated Myo1 phosphorylation is necessary to promote endocytic patch internalization, we examined endocytic patch behavior in the single point TEDS site phospho-null mutant, *myo1-S361A*, which cannot be phosphorylated by Pak1 but retains its C-terminal Arp2/3 binding domain (Attanapola et al., 2009). In the absence of Pak1-mediated phosphorylation of Myo1, we hypothesized that Fim1-mCherry patches would show similar weak internalization as observed in *myo1Δ* mutants.

In the TEDS site *myo1-S361A* mutant, we find that Fim1-mCherry is adequately recruited to endocytic patches, yet the patches do not properly internalize (Fig. 6 A and D, Fig. S2 A and B). Additionally, we observe that Myo1-S361A-GFP intensity remains mostly stable at the plasma membrane of polarized cell ends, while in *myo1^+^* cells, Myo1-GFP intensity increases as Fim1-mCherry is recruited to endocytic patches and is lost as Fim1-mCherry patches internalize at the cell ends (Fig. S2), as reported previously (Sirotkin et al., 2005). We also observe increased cytoplasmic intensity of Myo1-S361A-GFP compared to controls (Fig. S2 A and B). In *myo1-S361A* mutants, Fim1-mCherry lifetimes are indistinguishable from *myo1^+^* controls at the cell ends and sides, but patch lifetimes are prolonged at the division site (Fig. 6 C). Similar to *myo1Δ* mutants, the timing of Fim1 recruitment to endocytic patches is prolonged in *myo1-S361A* mutants (Fig. 6 D and H; Fig. S2). At the division site, Fim1 recruitment is enhanced in *myo1-S361A* mutants and peak Fim1-mCherry intensity is reached slightly later compared to *myo1^+^* controls (Fig. 6 I), mimicking *myo1Δ* mutants (Fig. 6 F). Similar to *myo1Δ* mutants, peak Fim1-mCherry intensity at the cell ends is reached later in *myo1-S361A* mutants compared to controls (Fig. 6 J). These observations suggest that branched actin formation can occur in the absence of Myo1 function, even though Myo1 is a nucleation promoting factor, albeit weak compared to Wsp1 (Sirotkin et al., 2005). We also find that while Myo1 regulates patch lifetimes at the cell sides, the TEDS site phosphorylation is not required for the patches at these sites. This aligns with the fact that the Pak1 kinase that phosphorylates the TEDS is also not required for patch dynamics at the cell sides.

### Cdc42 and Pak1 activity regulate the recruitment and dynamics of Myo1 at polarized sites

Our data show that Cdc42 and Pak1 are required for proper endocytic dynamics at polarized sites. We asked if Cdc42 and Pak1 regulate endocytosis via the regulation of Myo1 recruitment and dynamics at these sites. To test this, we first observed whether loss or delay of Cdc42 or Pak1 activation at the division site delays Myo1 recruitment to this site. Using *gef1Δ* mutants to delay Cdc42 activation at the division site as previously described (Fig. 2), we observe that Myo1-GFP recruitment to the division site is consequently delayed by ~10 mins compared to *gef1^+^* control cells (Fig. 7 A and B). Similarly, in *orb2-34* mutants lacking Pak1 kinase activity, Myo1-GFP recruitment to the division plane is delayed by ~4 mins compared to *orb2^+^* controls, even under permissive conditions (Fig. 7 C and D). These findings indicate that both Cdc42 and Pak1 activation promote timely recruitment of Myo1 to the division site.

**Figure 7.**
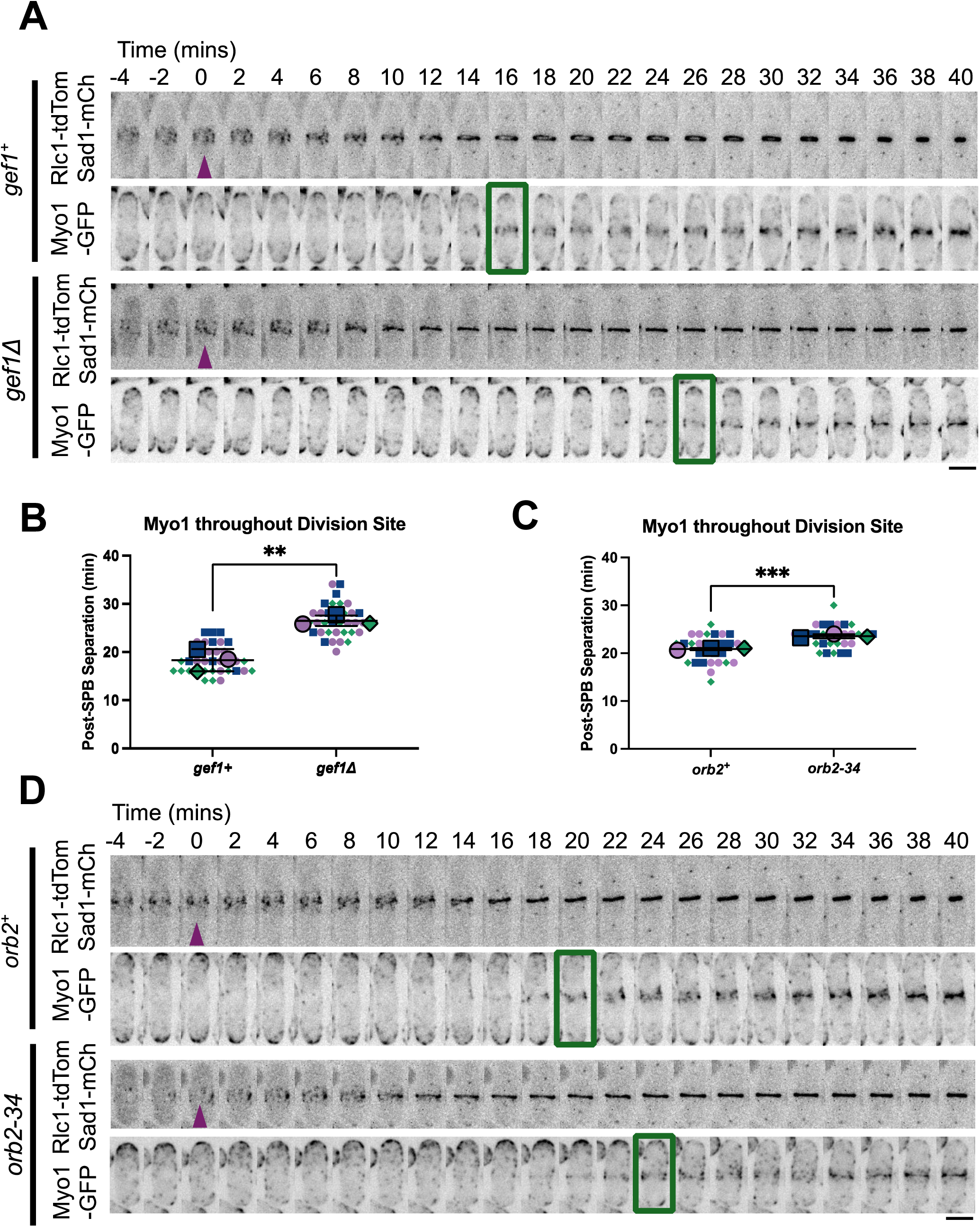
Myo1 localization to the division site requires Cdc42 and Pak1/Orb2 kinase. **A** and **D.** Representative cells showing Myo1-GFP appearance (green box) at the division site relative to Sad1-mCherry to mark spindle pole body (SPB) separation (purple arrowhead) in the indicated strains. **B** and **C.** Quantification of A and D, respectively, where time 0 = SPB separation. (n ≥ 10 cells per genotype per experiment). Unpaired Student’s *t*-test. ***, *p*<0.001; **, *p*<0.01, scale bar=5μm

Following our findings at the division site, we then asked if Cdc42 and Pak1 activation similarly regulate Myo1 recruitment to polarized cell ends. To test this, we first measured the mean intensity of Myo1-GFP recruited to the cell ends in genotypes with varying degrees of Cdc42 activity. While loss of *gef1* decreases Cdc42 activity (Wei et al., 2016), loss of the GAPs *rga4* and *rga6* increases Cdc42 activation levels (Campbell et al., 2022). Thus, we measured mean Myo1-GFP intensity at the cell ends of *gef1Δ* and *rga4Δrga6Δ* mutants compared to control cells. Intriguingly, Myo1-GFP recruitment is decreased in *gef1Δ* mutants and conversely enhanced in *rga4Δrga6Δ* mutants compared to controls (Fig. 8 A and B). To investigate the dynamics of Myo1 at the cell ends, we also performed 1 sec-interval time-lapse imaging and measured the dwell time of Myo1-GFP at the plasma membrane. These analyses reveal that Myo1-GFP dwell time is increased in both *gef1Δ* and *rga4Δrga6Δ* mutants compared to controls, suggesting that the myosin is less dynamic in both cases (Fig. 8 C and D).

**Figure 8.**
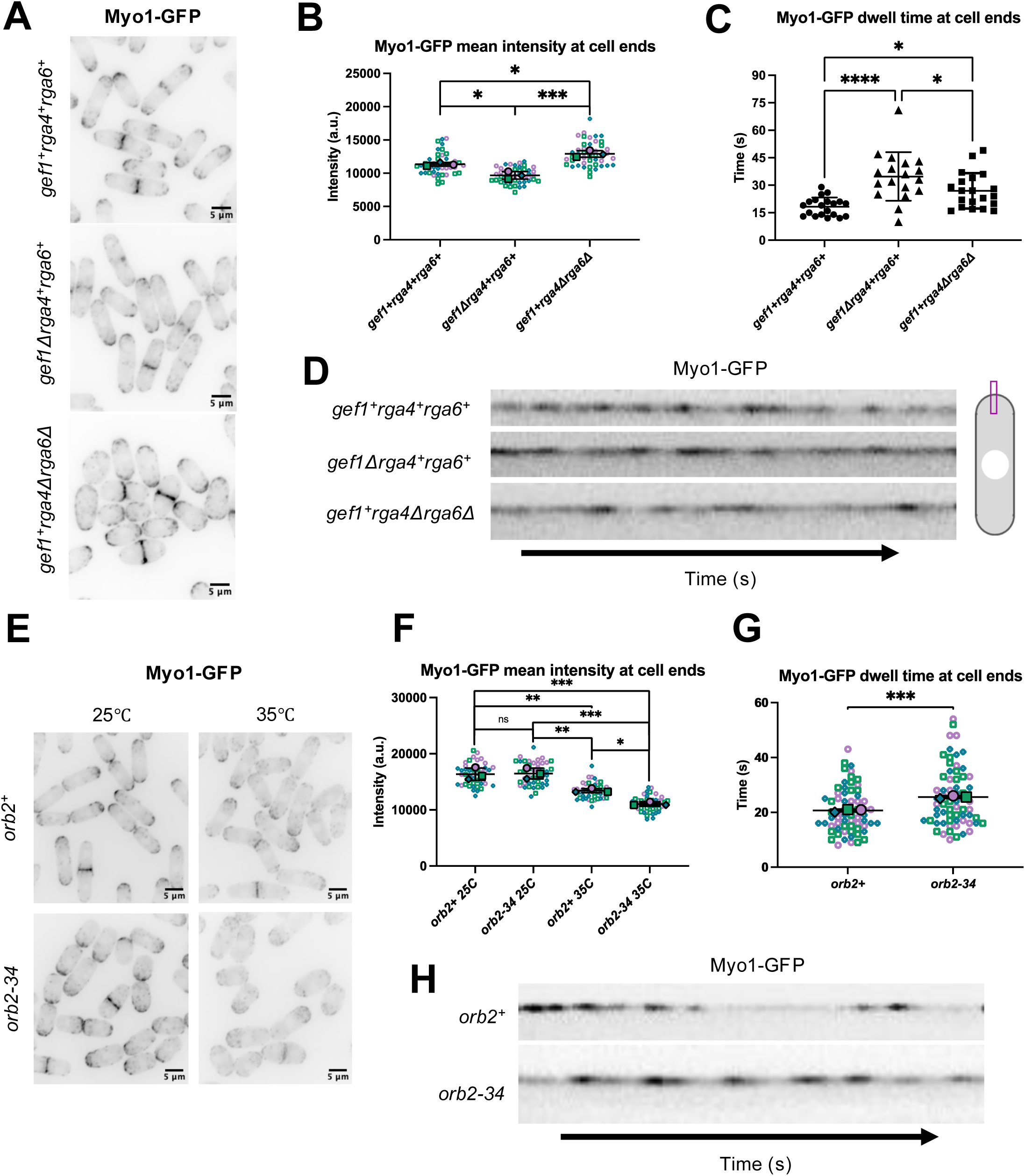
Cdc42 and Pak1/Orb2 kinase promotes Myo1 localization to the cell ends and division site. **A** and **E.** Sum Z-projections of Myo1-GFP in the indicated strains and conditions. **B** and **F.** Mean Myo1-GFP intensity measured at cell ends in the indicated strains (n=15 cells per genotype per triplicate experiment). **C** and **G.** Quantification of dwell times of Myo1-GFP in indicated strains. (n≥15 dwell time events per genotype. **D** and **H.** Montages showing Myo1-GFP dwell time dynamics at individual cell ends. One-way ANOVA with Tukey’s multiple comparisons post hoc test. ns, not significant; *, *p*<0.05; **, p.<0.01; ***, *p*<0.001; ****, p<0.0001, scale bar=5μm

We then performed similar experiments in *orb2-34* cells to see if decreased Pak1 activation impairs Myo1-GFP recruitment and dynamics at cell ends. Under permissive conditions at 25C°, mean Myo1-GFP intensity is indistinguishable between *orb2-34* and *orb2^+^* cell ends (Fig. 8 E and F). In contrast, temperature restriction at 35C° reduces mean Myo1-GFP intensity at the cell ends in both *orb2^+^* and *orb2-34* cells, although Myo1-GFP recruitment is more severely impacted in *orb2-34* mutants (Fig. 8 E and F). Finally, loss of Pak1 kinase function likewise impairs Myo1-GFP dynamics at cell ends, as Myo1-GFP dwell times at the plasma membrane are prolonged in *orb2-34* compared to *orb2^+^* cells under permissive conditions (Fig. 8 G and H). Together, these data indicate that Cdc42 and Pak1 activation promote the recruitment of Myo1 to the plasma membrane in a dose-dependent manner. Additionally, these results suggest that an intermediate range of Cdc42 activation facilitates proper Myo1 dynamics. In summary, our findings suggest that sufficient recruitment of Myo1 to polarized sites depends on both Cdc42 and Pak1 activation.

## DISCUSSION

In many cell-walled organisms, endocytosis is an actin-dependent process since cells must overcome high internal turgor pressure to properly bend and invaginate the plasma membrane (Basu et al., 2014; Lacy et al., 2018). In yeast, branched actin polymerization and crosslinking provide the force needed to promote successful patch internalization against high turgor pressure (Basu et al., 2014; Kaksonen and Roux, 2018; Skau et al., 2011). However, it is not known whether the force requirements are the same at distinct sites of endocytosis. For instance, at the division site, patches form in close proximity to the actomyosin ring, the synthesizing septum, as well as other membrane-associated cytoskeletal structures such as the septin ring (Wu et al., 2003), which may impact the force required for successful internalization. Here, we demonstrate that endocytic patch behavior is intricately linked with cell polarization. We find that endocytic patches at non-growing cell sides weakly recruit Fim1, display decreased lifetimes, and exhibit diffusive behavior. In contrast, endocytic patches within polarized regions recruit twice as much Fim1 with longer lifetimes, and exhibit directed motion during patch internalization. While it is unclear if the endocytic patches at the cell sides successfully undergo vesicle internalization. our data show that only the patches at the polarized sites exhibit the controlled and directed motion required for orderly membrane invagination and vesicle scission.

### Cdc42 and Pak1 activity promote initiation of endocytosis at sites of polarization

We show that Cdc42 activation is required for the initiation of branched actin-mediated endocytosis at the division site. Similar to Cdc42 activation patterns, Fim1-mEGFP first appears at the division site of *gef1^+^* controls during ring maturation well before the onset of ring constriction (Wang et al., 2016). However, in *gef1Δ* mutants, Fim1-mEGFP appearance is delayed to just before the onset of ring constriction, as reported for Cdc42 activation (Wei et al., 2016). Yet, once Cdc42 is finally activated in *gef1Δ* mutants, Fim1-mEGFP rapidly appears at the division site. These results suggest that the timing of Cdc42 activation and onset of endocytosis are tightly linked. In a similar manner, we also demonstrate that the Cdc42 downstream effector Pak1 promotes timely onset of endocytosis at the division site. Given the previously described challenges, we have not directly shown that Cdc42 activity is essential for endocytosis at the cell ends. However, we demonstrate that decreased Cdc42 activation via loss of its GEF *gef1* results in excessive Fim1 recruitment yet delayed patch internalization at the cell ends. Thus, Gef1-mediated Cdc42 activation facilitates proper patch formation and internalization at the cell ends.

### The two Cdc42 GEFs differentially regulate endocytosis

As we probed how Cdc42 activation promotes endocytosis, we found that various polarity mutants exhibit distinct phenotypes of endocytic patch behavior (summarized in Supplementary Table S1, S2, and S3). Individual loss of each Cdc42 GEF results in disparate phenotypes. Loss of *gef1 —*where *scd1* is the sole Cdc42 GEF— severely impacts endocytic patch behavior at polarized sites. On the other hand, loss of *scd1* —where *gef1* is the sole GEF— depolarizes the location of endocytic sites across the cell cortex but the patch dynamics remain mostly normal. These findings suggest that rather than strong, sustained activation via Scd1, transient, Gef1-mediated Cdc42 activation is required for proper endocytic patch formation and internalization. While dispensable for proper patch behavior, Scd1 is still required to prevent ectopic Cdc42 activation and thus regulates where endocytosis occurs. We observe that loss of *gef1* results in some differences between patch dynamics at the cell ends and the division site. In *gef1^+^* cells, peak Fim1-mEGFP patch intensity is reached ~2 seconds faster at the division site compared to the cell ends. In contrast, *gef1Δ* mutants appear to recruit Fim1 to endocytic patches at both sites with a delay. Additionally, Fim1-mEGFP patch lifetimes are increased at the cell ends of *gef1Δ* mutants. Comparatively, patch lifetimes are not enhanced at the division site of *gef1Δ* mutants, even though *gef1Δ* mutants recruit more Fim1 to endocytic patches. These observations indicate that the rate of branched actin synthesis is likely faster in *gef1Δ* mutants, since more branched actin is formed at the patch within the same span of time. As Cdc42 is under strong Scd1-mediated activation in *gef1Δ* mutants, it is possible that sustained Cdc42 activity in these mutants leads to enhanced branched actin synthesis. Our results further show that this excessive branched actin formation is actually deleterious to timely patch internalization in *gef1Δ* mutants. Together, these findings indicate that Cdc42 activation must be tightly regulated at the endocytic patch to form proper amounts of branched actin.

### Pak1 kinase differentially regulates endocytic patch dynamics at distinct sites

Although loss of *pak1* kinase function impairs endocytic patch internalization at the cell ends (Harrell et al., 2024), it seems that actin patch formation is most greatly impaired at the division site of *pak1* mutants. At restrictive temperature, Fim1 recruitment to endocytic patches at the cell ends is not impaired in *pak1* mutants, while it is greatly decreased at the division site. This indicates that patch assembly at the cell ends is more robust to loss of *pak1* function than the division site. However, as reported earlier, these patches do not internalize efficiently and often fail endocytosis (Harrell et al., 2024). This suggests that Pak1 could play additional roles in endocytosis apart from patch assembly. Alternatively, while the patch synthesizes sufficient actin filaments, it may not be properly organized for successful internalization. We also see enhanced patch assembly defects at the division site in *pak1* mutants. This suggests that Pak1 may play a more important role in at the division site that affects endocytosis. Already some such roles are known since Pak1 phosphorylates several cytokinetic ring proteins (Magliozzi et al., 2020).

### Cdc42 and Pak1 kinase regulate Type I myosin recruitment and dynamics at sites of endocytosis

We show that Cdc42 promotes Myo1 recruitment to polarized sites in a dose-dependent manner. Our data indicate that both insufficient and excessive Cdc42 activation leads to greater stability of Myo1 at the plasma membrane, as demonstrated by its increased dwell time in mutants of Cdc42 GEFs and GAPs and as well as the *pak1* kinase. This increased stability is reminiscent of the stable localization displayed by Myo1-S361A-GFP, which cannot be phosphorylated by Pak1 kinase at its motor domain TEDS site. Since endocytic patch internalization defects seem to coincide with increased Myo1 stability, these results suggest that Myo1 requires a certain level of dynamic behavior to properly promote endocytic patch internalization. Since Myo1 is a direct target of Pak1 (Attanapola et al., 2009), we initially expected *myo1-S361A* mutants to phenocopy the endocytic defects observed in kinase-dead *pak1* mutants. Our results, however, show distinct phenotypes of altered endocytic patch behavior for these mutants, which indicates that Pak1 has additional roles that are yet to be explored.

In line with previous reports, our findings suggest that the Arp2/3 complex can generate branched actin without Pak1 phosphorylation of Myo1’s TEDS site (Attanapola et al., 2009) and even without Myo1 altogether. Thus, the internalization defect observed in *myo1-S361A* mutants does not stem from an inability to create branched actin. Given that TEDS site phosphorylation promotes motor function of the Type I myosin in other organisms (Bement and Mooseker, 1995; Fujita-Becker et al., 2005; Pedersen et al., 2023), we suspect that the same likely occurs in *S. pombe*. Indeed, the perturbed localization and increased stability we observe for Myo1-S361A-GFP suggests that its motor function may be impaired. Such findings have been reported in budding yeast where Type I myosin motor activity is independent of its ability to nucleate actin (Manenschijn et al., 2019). As the other actin nucleation promoting factor, the WASP homolog Wsp1, activates Arp2/3 independently and more strongly than Myo1 in *S. pombe* (Sirotkin et al., 2005), we suspect that Wsp1 primarily promotes Arp2/3-dependent branched actin formation in *myo1-S361A* mutants.

Independent of Cdc42 and Pak1, Myo1 promotes patch assembly and dynamics at the cell sides. In agreement, Pak1-dependent TEDS site phosphorylation of Myo1 does not appear to be required at the cell sides. The patches at the cell sides contain less actin, have decreased lifetimes, and do not rapidly internalization into the cell interior. It is possible that while Myo1 enables actin patch assembly at the cell sides, it may not generate sufficient branched actin required for patch internalization. Sufficient branched actin assembly likely requires Pak1-dependent phosphorylation of Myo1. This would explain why the patches at the polarized sites show stronger internalization.

### Endocytic patch assembly and dynamics are differentially regulated at distinct cell sites

While both the growing cell ends and the division site are sites of polarization, we observe subtle differences in endocytic patch behavior between these sites. As endocytosis is a force-sensitive and actin-dependent process, the differences in patch dynamics at these distinct sites suggest that the forces experienced by patches at the division site are higher than that of the cell ends. If such is the case, patches at the division site would require stronger actin architecture and/or additional assistance from endocytic proteins such as Myo1 to overcome this increased force. This disparate force requirement could arise in several ways. Perhaps the flow of polarized secretion produces some counteractive force that must be overcome for patches to internalize. At the division site, polarized secretion occurs throughout the division plane, while endocytosis is largely confined to the outer rim of the membrane furrow (Wang et al., 2016). Furthermore, our observations that endocytic patches at the cell sides are weakly formed suggest that the forces they experience are also different than those at polarized regions. It is unclear if the patches at the cell sides internalize, but do not move as far from the plasma membrane since the force produced within the patch is less. It is possible that these patches do not require access to active Cdc42 to internalize. Alternatively, it is possible that they are random assemblies of branched actin that do not represent true sites of endocytosis.

Finally, it seems from our observations in *gef1Δ* mutants that formation of branched actin patches is not sufficient in itself to stimulate timely patch internalization, since patches excessively recruit both Cdc15 and Fim1 yet do not timely internalize. Perhaps the actin architecture within these mutants is improperly organized to internalize the patch. This hypothesis fits with our observations in *myo1Δ* and *myo1-S361A* mutants, which also assemble branched actin patches with excessive Fim1 yet exhibit severe patch internalization defects. Together, our findings indicate a direct role for Cdc42 in regulating endocytic patch formation via the Pak1 kinase and endocytic patch internalization via the Pak1 target, the Type I myosin, in a site-specific manner. A recent report characterized endocytic patch dynamics in different fungal species including *S. pombe* (Picco et al., 2024). Compared to the other species, endocytic patches in *S. pombe* appear to rapidly internalize at very high speeds deep into the cell interior (Picco et al., 2024; Sun et al., 2019), which suggests the patches may experience higher amounts of force which propel them inward at high velocity. Further investigation will determine if Cdc42, Pak1 and Myo1 differentially regulate endocytosis in different fungal species resulting in distinct dynamics.

## MATERIALS AND METHODS

### Strains and cell culture

The *S. pombe* strains used in this study are listed in Table 1. All strains are isogenic to the original strain PN567. Cells were cultured in yeast extract (YE) medium and grown exponentially at 25°C for 3 rounds of 8 generations before imaging. Standard techniques were used for genetic manipulation and analysis (Moreno et al., 1991).

### Microscopy

Imaging was performed at room temperature (23–25°C) for all experiments except for restrictive temperature experiments, which were performed at 35°C (Fig. 5). Images were acquired on a spinning disk confocal system equipped with a Nikon Eclipse inverted microscope with a 100×/1.49 NA lens, a CSU-22 spinning disk system (Yokogawa Electric Corporation), and a Photometrics EM-CCD camera (Photometrics Technology Evolve with excelon Serial No: A13B107000). On this system, images were acquired with MetaMorph (Molecular Devices, Sunnyvale, CA). Imaging was also performed on a 3i spinning disk confocal system using a Zeiss AxioObserver microscope equipped with a 100×/1.49 NA objective, an integrated Yokogawa spinning disk (Yokogawa CSU-X1 A1 spinning disk scanner), and a Teledyne Photometrics Prime 95b back-illuminated sCMOS camera (Serial No: A20D203014). On this system, images were acquired using SlideBook (3i Intelligent Imaging innovations). All images were analyzed with FIJI (National Institutes of Health, Bethesda, MD (Schneider et al., 2012)).

For still imaging, the cells were mounted directly on glass slides with a #1.5 coverslip (Thermo Fisher Scientific, Waltham, MA) and imaged immediately, and with fresh slides prepared every 10 min. *Z*-series images were acquired with a depth interval of 0.4 μm. For time-lapse images, cells were placed in 5 mm glass-bottom culture dishes (MatTek, Ashland, MA) and overlaid with YE medium containing 0.6% agarose and 100 μM ascorbic acid as an antioxidant to minimize toxicity to the cell, as reported previously (Frigault et al., 2009; Wei et al., 2017).

### Analysis of vesicle tracking

Wild-type and mutant cells expressing Fim1-mEGFP or Fim1-mCherry were grown to OD 0.2-0.5 and placed in glass-bottom culture dishes as previously described for time-lapse imaging. Cells were imaged in a single medial plane with a frame rate of 1 sec for ~3-5 minutes using the same laser power and exposure settings for all experiments. To measure patch lifetimes and mean fluorescence intensity of patches throughout their lifetimes, Fim1-mEGFP/Fim1-mCherry patches were tracked using the FIJI plugin TrackMate (Tinevez et al., 2017). Background fluorescence was subtracted from the imaging field away from cells to sharpen the signal of patches for ease of tracking. Lifetime was measured as the time from which a patch first displayed distinguishable fluorescence at the cell cortex to when the fluorescence was no longer detectable. Mean speed (μm/sec), MSD (mean squared displacement), and diffusion coefficients (μm^2^/sec) were all calculated using TrackMateR, an R package that can analyze data captured and saved by TrackMate as XML files ((Sittewelle and Royle, 2024); https://quantixed.org/2022/09/05/tracking-announcing-new-r-package-trackmater/). To ensure that patches were tracked from the beginning of patch formation through patch disassembly, patches selected for tracking 1. gradually increased in intensity until a peak intensity was reached and 2. decreased in intensity following peak intensity (Berro and Pollard, 2014; Wang et al., 2016). Patches tracked with TrackMate were also manually reviewed to ensure tracking accuracy.

### Disruption of Pak1 kinase function via temperature-sensitive allele *orb2-34*

Pak1 kinase was inactivated via the *orb2-34* (*pak1-ts*) temperature-sensitive mutation. Cells were grown for 2 days in YE medium. On the day of the experiment, cells were cultured to an OD of 0.2. A subset of these cells was incubated at the permissive temperature of 25°C, while the other set was incubated at the restrictive temperature of 35°C for 4 h. Cells were imaged after 4 h under temperature-controlled conditions.

### Analysis of Myo1-GFP mean fluorescence intensity at cell ends

Wild-type and mutant cells expressing Myo1-GFP were grown to OD 0.2-0.5 and imaged on slides. Still *Z*-images with 21-24 *z*-planes were collected at a *z*-interval of 0.4 µm for the 488 nm channel. The same number of *z*-slices were collected for wild-type and mutant cells for each experiment using the same imaging settings. FIJI was used to generate sum projections from the *z*-series and the polygon selection tool was used to encircle the “cap” of Myo1-GFP signal present at individual cell ends. Mean fluorescence intensity at individual cell ends was then measured within these manually selected Myo1-GFP “caps”. The background fluorescence in a cell-free region of the image was subtracted to normalize intensity measurements. Only cells found within the central 80% of the imaging field were used for analysis. This was done to account for uneven illumination of the imaging field.

### Analysis of Myo1-GFP dwell time at cell ends

Wild-type and mutant cells expressing Myo1-GFP were grown to OD 0.2-0.5 and placed in glass-bottom dishes as previously described for time-lapse imaging. Cells were imaged in a single medial plane with a frame rate of 1 sec for 3 minutes using the same imaging settings for all experiments. To visualize Myo1-GFP dwell time throughout the 3-minute time-lapse, kymographs were generated from 1 pixel-wide selection lines drawn across the midline of the cell end, parallel to the cell’s long axis. These kymographs were then manually analyzed to measure the dwell time of Myo1-GFP at individual cell ends in the following manner: Using FIJI’s rectangle selection tool, individual Myo1-GFP dwell time events were marked on each kymograph and the length of each event was measured in pixels. Since our imaging settings generated kymographs were 1 pixel = 1 sec, the number of pixels measured for an individual event equals Myo1-GFP dwell time in seconds for that event.

### Statistical tests

GraphPad Prism was used to determine significance. One-way ANOVA, followed by a Tukey’s multiple comparisons post-hoc test, was used to determine individual *p*-values when comparing three or more samples. When comparing two samples, an unpaired Student’s *t*-test (two-tailed, unequal variance) was used to determine significance.

## ACKNOWLEDGMENTS

We thank Drs. Vladimir Sirotkin and Daniel Mulvihill for strains; and Bret Judson for the Boston College imaging core facility.

## AUTHOR CONTRIBUTIONS

M.E.D. and B.F.C. conceived the project. M.E.D. obtained funding for the project. B.F.C., A.R.W., and U.P. performed experiments and analyzed data. B.F.C. and M.E.D. wrote the manuscript.

## DATA AND RESOURCE AVAILABILITY

Data and resource availability: All relevant data and resource can be found within the article and its supplementary information.

