## Supplemental Material for "Endocytic Patch Dynamics are Differentially Regulated at Distinct Cell Sites in Fission Yeast"

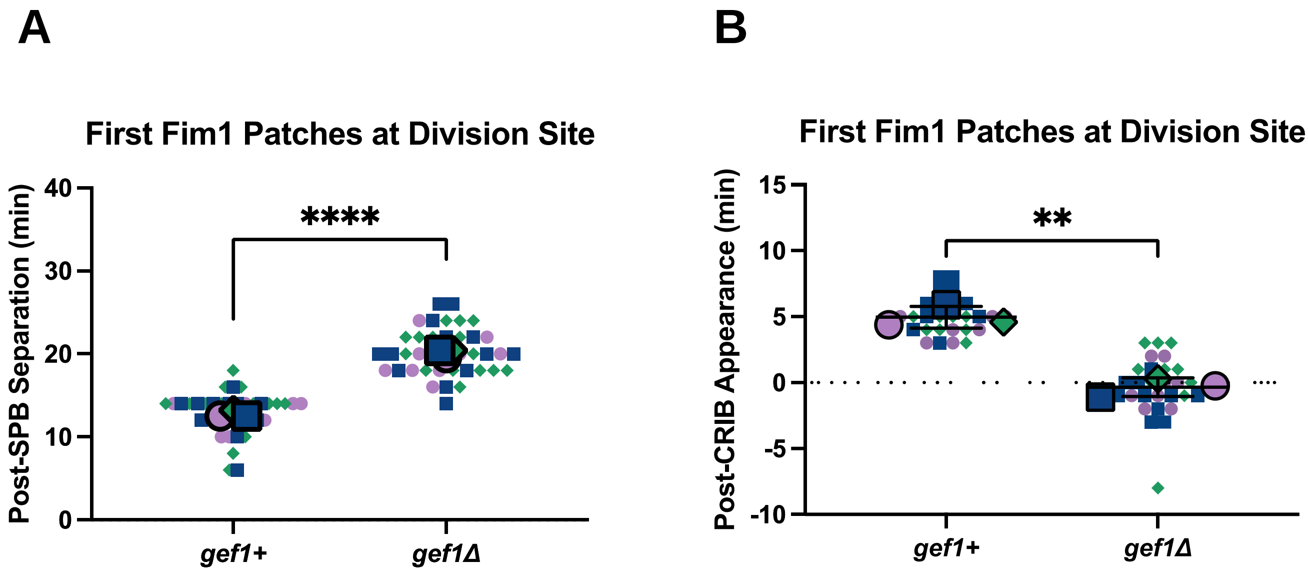


**Figure S1. Timing of initial appearance of Fim1 patches at the division site.** (A) Initial Fim1-mCh appearance relative to spindle pole body (SPB) separation. (B) Initial Fim1-mCh appearance relative to CRIB-3xGFP appearance at the division site. (n ≥ 8 cells per genotype per experiment). Unpaired Student's *t*-test. ****, $p < 0.0001$; **, $p < 0.01$

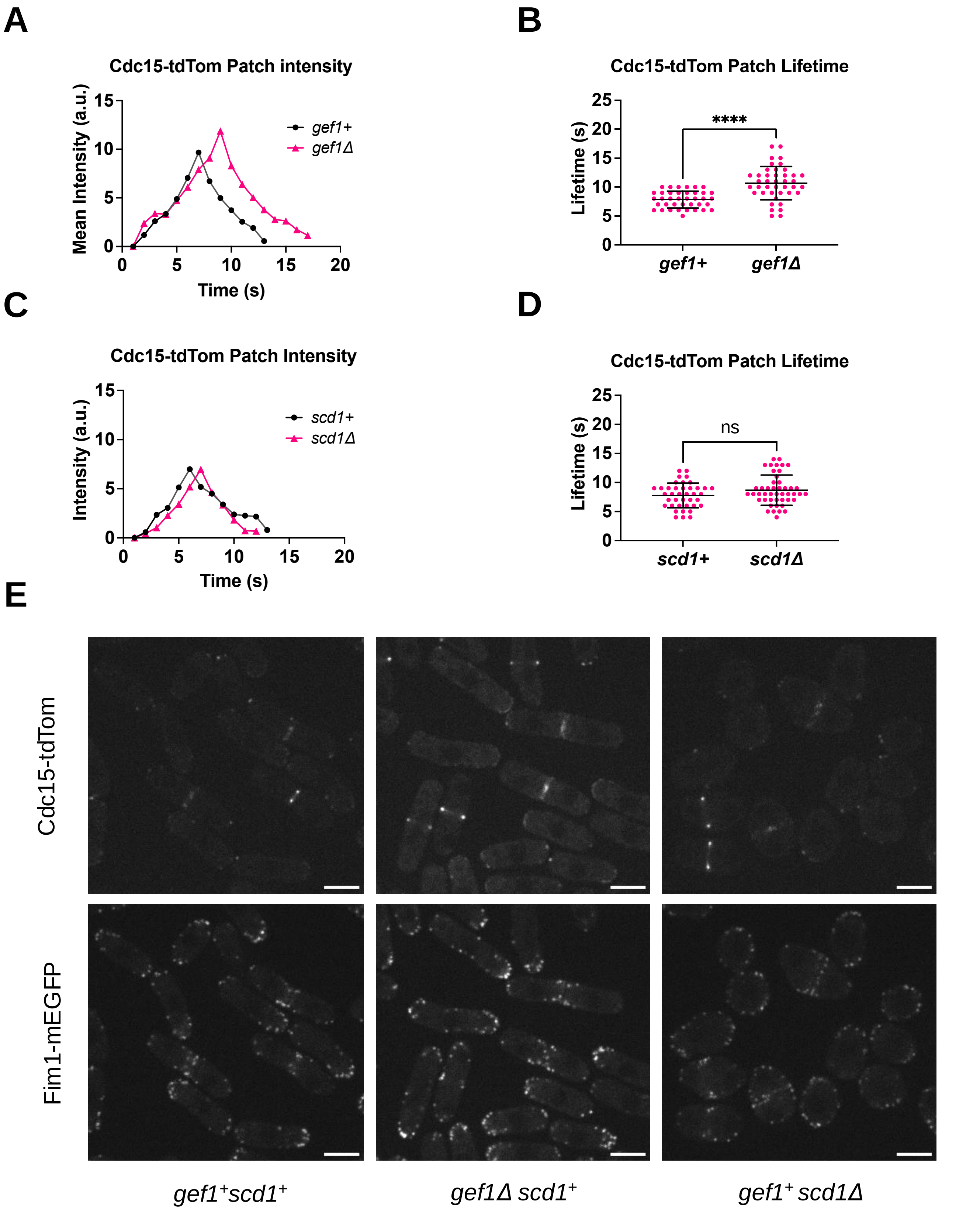


**Figure S2. Loss of Gef1 increases the accumulation of Cdc15 within endocytic patches.** (A) and (C) Mean patch intensity of Cdc15-tdTom at the cell ends in the indicated genotypes. (B) and (D) Cdc15-tdTom patch lifetime at the cell ends in the indicated genotypes (n ≥ 32 patches per genotype. Unpaired Student's *t*-test. ****, $p<0.0001$; ns, not significant

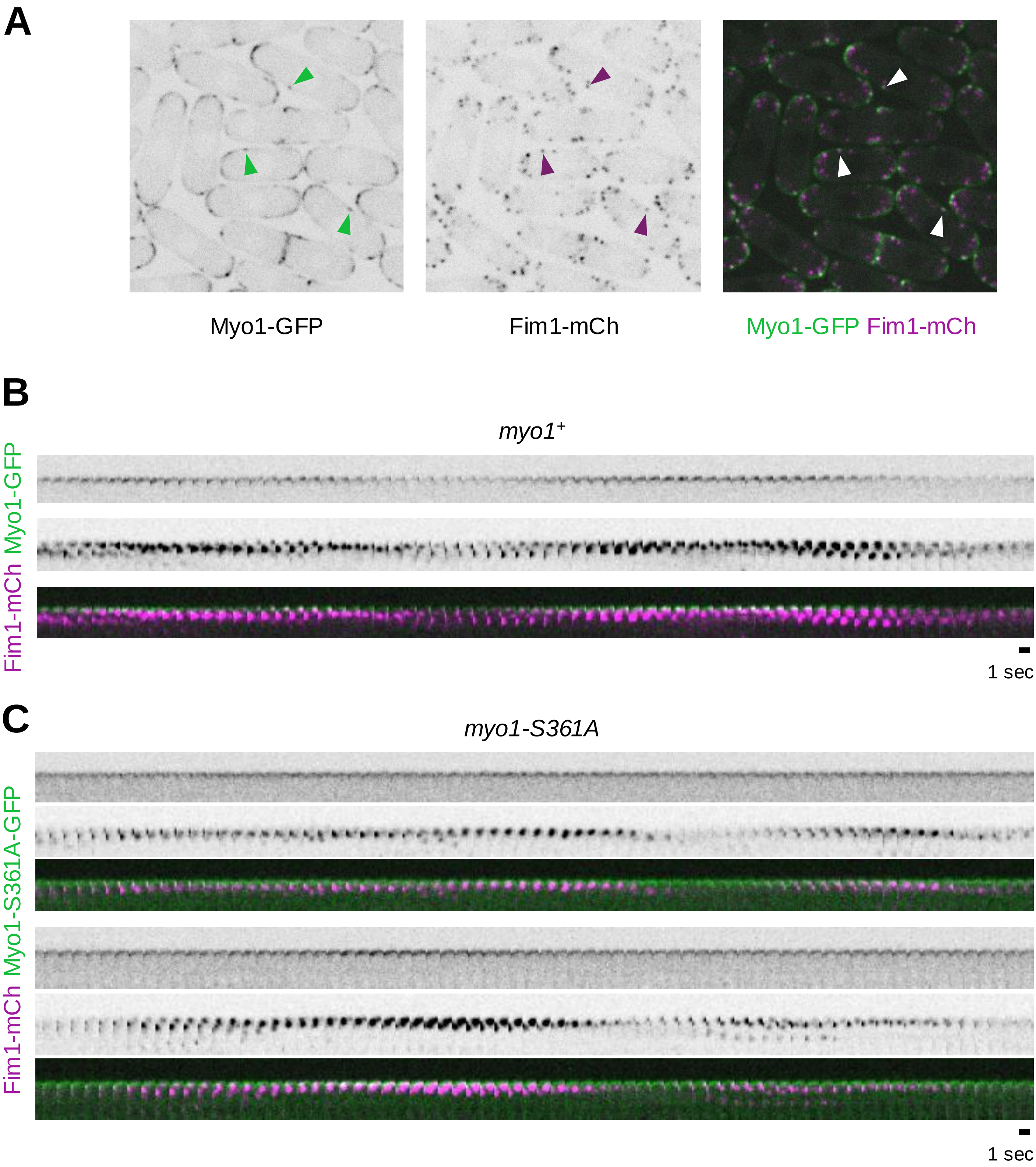


**Figure S3. Myo1 promotes Fim1 internalization at the polarized ends and the division site, yet not at the cell sides.** (A) Wild-type cells expressing Myo1-GFP and Fim1-mCh show that both Myo1 and Fim1 localize to the cell sides as well as to polarized cell ends and the division site. **(**B) and (C) Montages of Fim1-mCh and Myo1-GFP at the cell ends of the indicated genotypes (scale bar = 1 sec).

| **Cell ends** | **Fim1 recruitment rate** | **Fim1 peak recruitment** | **Fim1 lifetime** | **Timing of patch internalization** |
|---|---|---|---|---|
| control | normal | normal | normal | normal |
| *gef1Δ* | normal | ↑ | ↑ | delayed |
| *scd1Δ* | normal | normal | ↓ | normal |
| *myo1Δ* | ↓ | ↑ | ↑↑ | delayed |
| *myo1-S361A* | semi-normal | normal | normal | delayed |

**Table. S1.** Phenotypes of Fim1-mEGP or Fim1-mCh endocytic patch behaviors at the cell ends in various polarity mutants compared to controls.

| **Division site** | **Fim1 recruitment rate** | **Fim1 peak recruitment** | **Fim1 lifetime** | **Timing of patch internalization** |
|---|---|---|---|---|
| control | normal | normal | normal | normal |
| *gef1Δ* | ↓ | ↑ | normal | delayed |
| *scd1Δ* | normal | normal | ↓ | normal |
| *myo1Δ* | ↓↓ | ↑↑ | ↑↑ | very delayed |
| *myo1-S361A* | semi-normal | ↑ | ↑ | delayed |

**Table S2.** Phenotypes of Fim1-mEGP or Fim1-mCh endocytic patch behaviors at the division site in various polarity mutants compared to controls.

| **Cell ends** | **Fim1 recruitment rate** | **Fim1 peak recruitment** | **Fim1 lifetime** | **Timing of patch internalization** |
|---|---|---|---|---|
| 25ºC control | normal | normal | normal | normal |
| 25ºC *orb2-34* | normal | ↓ | normal | normal |
| 35ºC control | ↓ | ↓ | ↓ | early |
| 35ºC *orb2-34* | semi-normal | same as *orb2-34* at 25ºC | ↓ | early |
| **Division site** | | | | |
| 25ºC control | normal | normal | normal | normal |
| 25ºC *orb2-34* | normal | ↓ | ↓ | normal |
| 35ºC control | semi-normal | normal | ↓ | early |
| 35ºC *orb2-34* | ↓↓ | ↓↓ | same as control at 35ºC; similar to *orb2-34* at 25ºC | same as control at 35ºC |

**Table S3.** Phenotypes of Fim1-mEGP endocytic patch behaviors in *orb2-34* mutants compared to *orb2*$^{+}$ controls at permissive (25°C) and restrictive (35C°) temperatures.

| Strain | Genotype | Source |
|---|---|---|
| YMD 2098 | *fim1-mEGFP-kanMX cdc15-tdTom-NAT ade6- leu1-32 ura4-d18* | This study |
| YMD 2104 | *gef1Δ::ura4*$^+$ *fim1-mEGFP-kanMX cdc15-tdTom-NAT ade6- leu1-32 ura4-d18* | This study |
| YMD 2159 | *scd1Δ::ura4*$^+$ *fim1-mEGFP-kanMX cdc15-tdTom-NAT ade6- leu1-32 ura4-d18* | This study |
| YMD 2084 | *fim1-mEGFP-kanMX rlc1-tdTom-NAT sad1-mCh-KanMX ade6- leu1-32 ura4-D18* | This study |
| YMD 2085 | *gef1Δ::ura4*$^+$ *fim1-mEGFP-kanMX rlc1-tdTom-NAT sad1-mCh-kanMX ade6- leu1-32 ura4-D18* | This study |
| YMD 2250 | *CRIB-3xGFP-ura4*$^+$ *fim1-mCh-NAT ade6- leu1-32 ura4-D18* | This study |
| YMD 2253 | *gef1Δ::ura4*$^+$ *CRIB-3xGFP-ura4*$^+$ *fim1-mCh-NAT ade6- leu1-32 ura4-D18* | This study |
| YMD 2497 | *orb2-34-ts fim1-mEGFP-kanMX rlc1-tdTom-NAT sad1-mCh-KanMX ade6- leu1-32 ura4-D18* | This study |
| YMD 1930 | *fim1-mEGFP-kanMX rlc1-tdTom-NAT ade6- leu1-32 ura4-D18* | This study |
| YMD 1937 | *orb2-34-ts fim1-mEGFP-kanMX rlc1-tdTom-NAT ade6- leu1-32 ura4-D18* | This study |
| VS 888-3 YMD 2064 | *fim1-mCh-NAT ade6- leu1-32 ura4-D18* | Vladimir Sirotkin |
| YMD 2324 | *myo1Δ::kanMX fim1-mCh-NAT ade6- leu1-32 ura4-D18* | This study |
| YMD 2074 | *myo1-GFP-KanMX fim1-mCh-NAT ade6- leu1-32 ura4-D18* | This study |
| YMD 2227 | *myo1Δ::kanMX leu1::nmt41-GFP-myo1-S361A fim1-mCh-NAT ade6- ura4-D18* | This study; made using *myo1-S361A* mutant from Attanapola et al., 2009 |
| YMD 2480 | *myo1-GFP-kanMX rlc1-tdTom-NAT sad1-mCh-kanMX ade6- leu1-32 ura4-D18* | This study |
| YMD 2481 | *gef1Δ::ura4*$^+$ *myo1-GFP-kanMX rlc1-tdTom-NAT sad1-mCh-kanMX ade6- leu1-32 ura4-D18*``` | This study |
| YMD 2496 | *orb2-34-ts myo1-GFP-kanMX rlc1-tdTom-NAT sad1-mCh-kanMX ade6- leu1-32 ura4-D18* | This study |
| YMD 734 | *myo1-GFP-kanMX rlc1-tdTom-NAT ade6- leu1-32 ura4-D18* | Lab stock |
| YMD 1807 | *rga4Δ::ura4*$^+$ *rga6Δ::kanMX myo1-GFP-kanMX rlc1-tdTom-NAT ade6- leu1-32 ura4-D18* | This study |
| YMD 1907 | *gef1Δ::ura4*$^+$ *myo1-GFP-kanMX rlc1-tdTom-NAT ade6- leu1-32 ura4-D18* | This study |
| YMD 1915 | *orb2-34-ts myo1-GFP-kanMX rlc1-tdTom-NAT ade6- leu1-32 ura4-D18* | This study |

**Table S4.** List of strains used in this study
